# Two-Step Small Scale Purification of Recombinant Adeno-Associated Viruses

**DOI:** 10.1101/715029

**Authors:** Shih-Heng Chen, Amy Papaneri, Mitzie Walker, Erica Scappini, Robert D. Keys, Negin P. Martin

## Abstract

Recombinant adeno-associated viruses (AAVs) are robust and versatile tools for *in vivo* gene delivery. Natural and designer capsid variations in AAVs allow for targeted gene delivery to specific cell types. Low immunogenicity and lack of pathogenesis also add to the popularity of this virus as an innocuous gene delivery vector for gene therapy. AAVs are routinely used to express recombinases, sensors, detectors, CRISPR-Cas9 components, or to simply overexpress a gene of interest for functional studies. High production demand has given rise to multiple platforms for production and purification of AAVs. However, most platforms rely heavily on large amounts of starting material and multiple purification steps to produce highly purified viral particles. Often, researchers require several small-scale purified AAVs. Here, we describe a simple and efficient technique for purification of recombinant AAVs from small amounts of starting material in a two-step purification method. In this method, AAVs are released into the packaging cell medium using high salt concentration and pelleted by ultracentrifugation to remove soluble impurities. Then, the resuspended pellet is purified using a protein spin-concentrator. The two-step purification consisting of ultracentrifugation and spin-concentration eliminates the need for fraction collection and the time-consuming evaluation of individual fractionated aliquots for titer and purity. In this method, the resulting AAV preparations are comparable in titer and purity to commercially available samples. This simplified process can be used to rapidly generate highly purified AAV particles in small scale, thereby saving resources.

## Introduction

Recombinant adeno-associated viruses (rAAVs, referred to as AAVs in this manuscript) are replication-defective parvoviruses that readily infect dividing and non-dividing mammalian cells [1]. AAVs can deliver up to 4.5 kilobase pairs (kbp) of genetic material that is often maintained epigenetically and therefore does not cause insertional mutation. In rare cases, the AAV genome is inserted at regions of host chromosomal instability and breakage. The AAV genome consists of a single stranded DNA with self-complementary ends that form high-molecular-weight head-to-tail circular concatemers [2]. These concatemeric circles are often maintained in transduced cells for lasting gene expression *in vivo* [3].

AAVs are promising agents for gene therapy due to their varying capsid tropism, low immunogenicity, and lasting gene expression [1,4]. In research, they are ideal for functional studies *in vivo*. The genetic load can accommodate constitutive or inducible promoters to regulate expression of genes of interest in addition to selectable and/or fluorescent markers. Many new techniques such as Cre-recombination-based AAV targeted evolution (CREATE) have given rise to a myriad of AAV capsids to achieve the desired performance and cell/tissue targeting [5–9]. Biotech manufacturing has responded to the demand for AAVs by increasing quality and quantity of produced AAV vectors while maintaining reasonable cost.

Currently, there are several platforms used for production and purification of AAVs [10,11]. Biotech manufacturing and virus core facilities often rely on helper viruses such as adenovirus, herpesvirus, or baculoviruses for *en mass* delivery of complementary genes for large-scale and cost-efficient AAV production. In traditional approaches, complementary genes are delivered as plasmids to eukaryotic cells with a variety of transfection reagents [12,13]. Human embryonic kidney 293 cells (HEK293 and HEK293T) are a preferred cell line for packaging AAVs since they constitutively express adenovirus E1a/b factors that are needed for packaging, are economical, and efficiently transfected [12]. To produce AAVs, eukaryotic cells are transfected with three plasmids: 1) an AAV transfer vector carrying the gene of interest flanked by Inverted Terminal Repeats (ITRs), 2) Rep/Cap genes, and 3) Helper plasmid delivering VA RNAs, E2A, and E4OEF6 genes [12]. Following transfection, AAVs are collected from the media or cell lysate and subjected to numerous purification steps [13–17]. Gradient centrifugation with cesium chloride (CsCl) or iodixanol provides flexibility for AAV purification since it can be used to purify different AAV serotypes. However, using gradients to purify AAV is time consuming and requires multiple purification steps to produce high purity AAV. The formed gradients are fractionated and individually evaluated for amount and purity of AAV particles. Moreover, CsCl can exert toxic effects in animals and therefore, dialysis of fractions containing AAV with a physiologically balanced solution is necessary before use *in vivo* [15,17,18]. As expected, AAV particles are lost after each purification step, and therefore, large amounts of starting material are needed to ensure sufficient amount of AAV recovery. Complexity of purification steps and the length of time often prevents small laboratories from preparing their own AAV samples. Therefore, customized AAV preparations are commercially packaged for a significant cost or research in labs with limited budgets is planned based on available AAV stock preparations.

Here, we describe a method for a small-scale production of AAVs that allows research laboratories to produce purified virus rapidly, efficiently, and economically. Using this method, we compared and analyzed yields for AAV preparations with serotypes 1, 2, 5, 6, 8, and 9. The process effectively isolated infectious AAV particles for all serotypes except for serotype 2. Reduced number of steps in the purification process minimized virus particle loss and saved time. This method allows for a simple, rapid, and efficient virus purification from a small number of transduced cells.

## Materials and Methods

### Animals

C57BL/6J mice were obtained from Jackson Laboratories (Bar Harbor, ME, USA). Mice were housed in polycarbonate cages in animal facilities with controlled environmental conditions with a 12-hour artificial light-dark cycle and were provided fresh deionized water and NIH 31 chow *ad libitum*. All animal procedures were approved by the Institutional Animal Care and Use Committee and conducted in strict accordance with the National Institutes of Health animal care and use guidelines.

### Cell culture

Mycoplasma-free HEK293-AAV cells (Cell Biolabs Inc., Cat. # AAV-100) were maintained in Dulbecco’s modified Eagle’s medium (DMEM, Life Technologies, Grand Island, NY, USA) supplemented with 10% fetal bovine serum (FBS, HyClone Laboratories, South Logan, UT, USA), 2 mM L-glutamine, and 1 mM sodium pyruvate. Cells were passaged three times per week to maintain them in exponential growth phase.

### Plasmids and viruses

The plasmids used for transfection were listed (1) cis plasmid pAAV.hSyn.eGFP.WPRE.bGH (Addgene Cat# 105539) and pAAV-hSyn-eGFP (Addgene Cat# 50465); (2) trans plasmids pAAV2/1 (Cell Biolabs Inc., Cat# VPK-421), pAAV2/2 (Cell Biolabs Inc., Cat# VPK-422), pAAV2/5 (Cell Biolabs Inc., Cat# VPK-425), pAAV2/6 (Cell Biolabs Inc., Cat# VPK-426), pAAV2/8 (Cell Biolabs Inc., Cat# VPK-428), or pAAV2/9 (UPenn Vector Core); (3) pHelper plasmid containing adenovirus E2A, E4 and VA genes (Cell Biolabs Inc., Part No. 340202). AAV2-hSyn-EGFP (Addgene Cat# 50465-AAV2) and AAV9-hSyn-EGFP (UPenn Vector Core Cat # AV-9-PV1696) were purchased from Addgene and University of Pennsylvania Vector Core, respectively, as positive controls in this study.

### AAV production and two-step purification

For transfections, each 15-cm dish was seeded with HEK293-AAV cells at 6 × 10^6^ cells in 20 ml of DMEM with 10% FBS without antibiotics. Cells were incubated at 37ºC in 5% CO_2_ for 24 hours before transfection. The AAV cis, AAV trans and pHelper plasmids (33.3 ug of each) were added to 2 ml of sterile 150 mM NaCl solution. Polyethylenimine MAX (PEI “MAX”, Polysciences, Warrington, PA, USA) stock solution was prepared at 16 mg/ml in sterile water and the pH was adjusted to 4.5 with sodium hydroxide. 12.5 µl of PEI stock was added to the 2 ml plasmid solution and mixed by vortexing. After 10 minutes of incubation at room temperature, 2 ml of solution was added dropwise to a 15-cm plate. Cultures were incubated at 37°C in 5% CO2 incubator for 24 hours and then the media was changed to 14 ml of fresh serum-free DMEM containing 2 mM L-glutamine, and 1 mM sodium pyruvate. 120 hours post-transfection, 2ml of 5M NaCl was added to the plates and incubation was resumed for an additional 2 hours before collecting the culture medium into 50ml conical tubes. Turbonuclease (Eton Bioscience, San Diego, CA, USA) was added to the culture supernatant to a final concentration of 50 units/ml and incubated at 37°C for 1 hour. To remove the cellular debris, the collected culture medium was centrifugated at 3,000 x g for 20 minutes at 4°C. After centrifugation, the supernatant was collected and filtered through a 0.45 µm Durapore PVDF filter (EMD Millipore, Billerica, MA, USA). Filtered supernatant was aliquoted into 30 ml conical tubes (Beckman Coulter, Brea, CA, USA) underlaid with a 4 ml sucrose cushion (40% sucrose in Tri-sodium chloride-EDTA buffer/TNE, sterile filtered) and centrifuged for 16 hours at 100,000 x g in a Beckman Coulter SW32Ti rotor at 4°C to pellet the AAV virus. After centrifugation, supernatant was gently removed, and the viral pellet was resuspended in 0.5 ml of cold PBS and mixed on a nutator at 4°C overnight. The next day, the virus was mixed by pipetting up and down in the tubes, combined, and then diluted with additional PBS to a final volume of 10 ml. Pooled virus solution was cleared of debris by centrifugation at 500 × g at 4°C for 10 minutes. The remaining supernatant was concentrated in a 100-kDa molecular weight cutoff (MWCO) protein concentrator (Pierce, Rockford, IL, USA) by centrifugation at 3,000 x g at room temperature for 10-minute intervals until the volume was reduced to 100-500 μl. The virus was aliquoted and stored at −80°C.

### AAV titer by Q-PCR

The AAV genome was titered as previously described [17]. Briefly, AAV samples were serially diluted from 10^−2^ to 10^−8^-fold, and the standard curve was generated by diluting the AAV plasmid containing the ITR2 sequence (from 10^5^ to 32 copies per 5 µl). The Q-PCR was performed using SYBR Green PCR Master Mix (Applied Biosystems, Foster City, CA, USA) and a LightCycler 96 System (Roche, Indianapolis, IN, USA) according to manufacturer’s protocols. The ITR forward primer (5’ GGAACCCCTAGTGATGGAGTT 3’) and the ITR reverse primer (5’ CGGCCTCAGTGAGCGA 3’) were added to the reaction with final concentration of 300 nM in 25 µl of total volume (5 µl of sample or standard curve plus 20 µl of primers and SYBR mix), and under the following PCR conditions: 95°C for 10 minutes, 95°C for 10s, 60°C for 60s, for 45 cycles. The virus titer was calculated by LightCycler 96 SW1.1 software using the parallel standard curve in the reaction, and the titer was given in genome copy/ml (GC/ml).

### Silver staining

The purity of AAV was determined via polyacrylamide gel electrophoresis (PAGE) by resolving 3-5 × 10^10^ GC of each AAV sample boiled in NuPage sample buffer, loaded onto a 4-12% NuPage Bis-Tris gels (Thermo Fisher Scientific, Rockford, IL, USA). The proteins were revealed using Pierce Silver Staining kit (Thermo Fisher Scientific, Rockford, IL, USA) according to manufacturer’s protocols. The image was acquired by GeneFlash Bio Imaging system (Syngene, Frederick, MD, USA).

### Organotypic mouse brain slice culture and AAV transduction

Brain slices were isolated from C57BL/6J mice (postnatal 6-8 days). Mice were humanely and rapidly sacrificed. Brain slices were prepared according to previously described methods [19–21]. 350 μm slices of brain were sectioned using a vibratome (Leica VT1200 S, Leica Biosystems, Buffalo Grove, IL, USA) in ice-cold cutting solution (GIBCO MEM Cat#: 61100, NaHCO3 26 mM, HEPES 25 mM, Tris 15 mM, Glucose 10 mM, MgCl_2_ 3 mM). The organotypic brain slices were placed on a 0.4-μm pore size membrane insert (Transwell 3450-Clear, Corning Incorporated, Corning, NY, USA) in a 6-well plate. Slices were cultured at 37°C and 5% CO_2_ with the medium changed three times per week. AAVs were added as a drop on top of brain slices at 24 hours post dissection and incubated for 2 weeks. To visualize cells infected with AAVs, fluorescence images were captured on a Zeiss LSM710 (Carl Zeiss Inc, Oberkochen, Germany) using an EC Plan-Neofluar 10x/0.3 objective. The 488 nm laser line from an Argon laser at 5% power was used for excitation of the GFP labeled cells; after which, a 493-577nm band pass emission filter was used to collect the images of the GFP signal.

### TEM electron microscopy

Transmission electron microscopy was performed on equal volumes (5 μL) of AAV9 samples that were either prepared by the two-step purification method or purchased from the UPenn Core. Samples were prepared for negative staining by dropping 5 μl aliquot of viral suspension on a formvar coated 300 mesh grid and blotting off the grid at the edge with Whatman’s #1 filter paper until not quite dry. The wet grid was immediately negative stained with a drop of 1% phosphotungstic acid, pH 7.1, for 1 minute, and blotted dry at the edge with Whatman’s #1 filter paper. Negative stained images were obtained at 68000x.

## Results

### Two-step purification of AAVs

In order to assess the efficiency and yield of the two-step purification protocol, the same AAV transfer vector, pAAV-hSyn-GFP, was packaged and purified using serotypes 1, 2, 5, 6, 8, and 9 as described in the Materials and Methods section. The purification steps from seeding the HEK293-AAV cells to aliquoting and freezing the samples were completed in 8 days (Fig. 1). Total amount of time per day spent on the purification process is significantly less than traditional purification methods since the need for fraction collection, assaying each fraction, and further concentration and dialysis to remove chemicals and impurities is eliminated. Polyethylenimine MAX (PEI) was used as an economical and efficient method to transfect and deliver packaging plasmids to HEK293-AAV cells [17]. Other transfection methods, including calcium phosphate precipitation of DNA, can be substituted for PEI transfection. Cells can be grown in attached monolayers, multilayered flasks, or in suspension depending on the preparation size and user preference. PEI offers the versatility of transfection in adherent or cell suspension cultures.

**Fig. 1.**
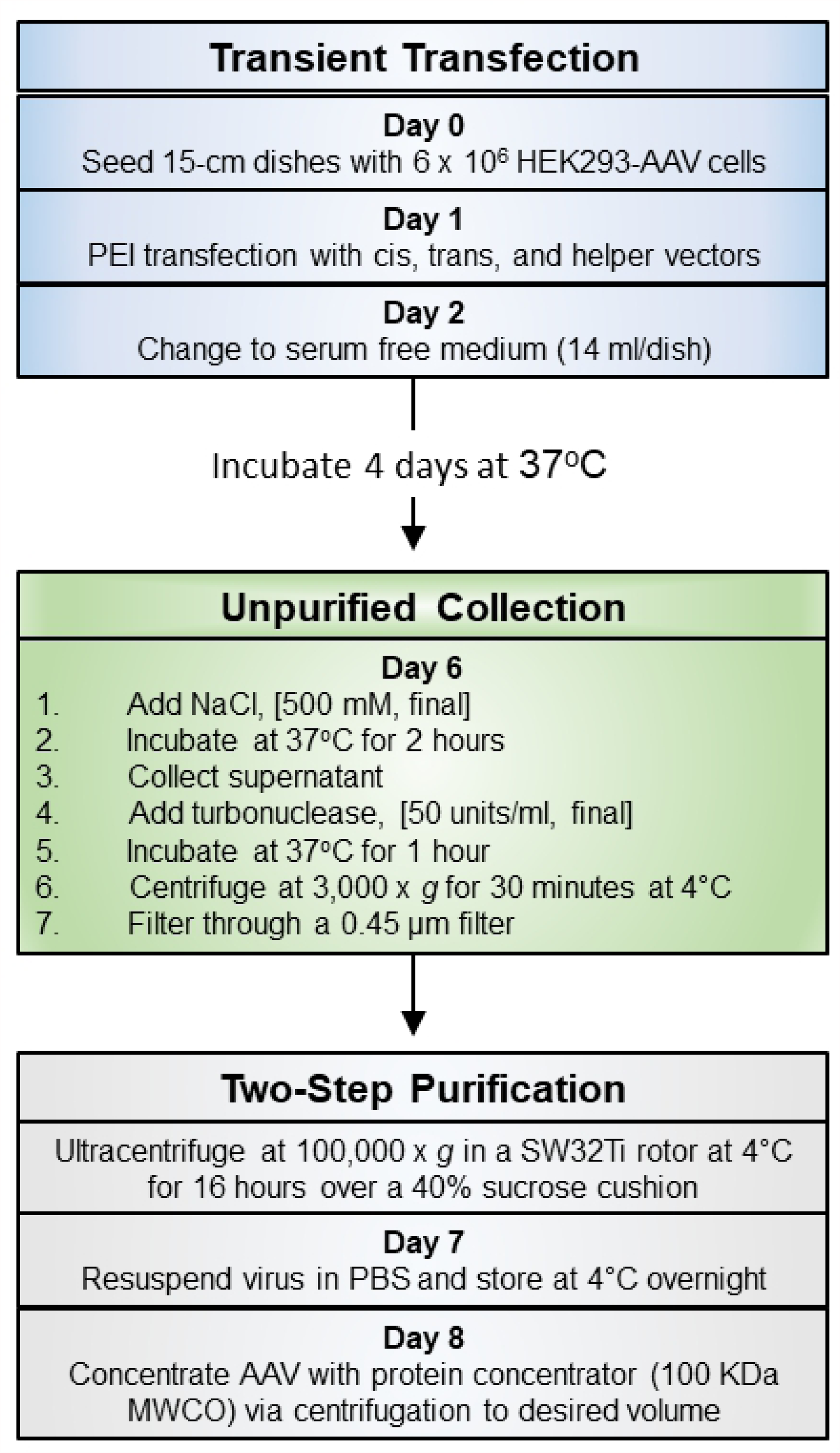
Protocol outline.

The NIEHS Viral Vector Core AAV preparation protocol outline, including the two-step AAV purification process. The entire procedure is completed in 8 days with less processing time per day during purification steps.

Transfected cells were allowed to express AAV packaging genes and incubated at 37°C for 5 days. Recent studies have shown that extending incubation time from 3 to 5 days after transfection results in release of AAV particles from cells and their accumulation in culture medium [14,17,22]. In order to minimize protein content in the cell medium, the media was switched to serum-free medium 24 hours post transfection. Incubation with high salt (500mM) has been shown to increase the release of AAV particles from the cells [17,23]. Salt’s effect on protein-protein interactions is often responsible for increased protein solubility [24]. Mature AAV particles are stable under high salt and in a wide range of pH conditions. Therefore, salt concentration was increased to enhance AAV particle release from cells without damage to the particles.

AAV particles were collected in supernatant media, cleared from debris, and treated with Turbonuclease to remove RNA, chromosomal DNA and the packaging plasmid remnants. At this stage, the unpurified AAV particles were suspended in a pool of protein impurities. In traditional purification methods, the unpurified AAV solution is subject to two rounds of gradient centrifugation on cesium chloride or a single centrifugation in a more inert medium such as iodixanol [17]. The collected fractions are then individually assayed for purity, pooled, and then further concentrated. In this two-step purification protocol, crude AAV particles are pelleted by ultracentrifugation over a 40% sucrose cushion. After ultracentrifugation, less than 15% of AAV particles (genocopies determined by Q-PCR) are discarded in the medium and the remainder of particles are pelleted below the sucrose cushion. The resulting pellet is then resuspended in phosphate buffered saline (PBS). Although many soluble AAV impurities are removed by pelleting the virus and do not precipitate below the 40% sucrose cushion, the resuspended AAV pellets still contain impurities that are subsequently removed by using a 100-kDa molecular weight cutoff (MWCO) protein concentrator. Approximately 10% of AAV genocopies (determined by Q-PCR) are lost through the protein concentrator pores during centrifugation. The volume of AAV solution can be continuously reduced by centrifugation until the desired volume is reached.

### AAV titers and purity

The number of AAV genocopies (GC) and the titer for individual preparations were determined by standard Q-PCR protocols as described in Material and Methods [17]. These titers were originated from small-scale AAV preparations from five or six 15-cm plates (Table 1). AAV serotypes 1, 5, 8, and 9 yielded titers of greater than 1E+12 genome copies per ml (GC/ml) in volumes greater than 100 μL per preparation. Greater than 1500 genome copies were recovered per HEK293 cell. AAV serotypes 2 and 6 had low yields in similar volumes. AAV with serotype 8 produced the greatest yield of 1.72E+05 genome copies per cell (Table 1).

**Table 1.** Virus yield of AAV serotypes using two-step purification method

| AAV serotype | Transgene | Virus titer (GC/ml) | Final volume (μl) | Total virus yield (GC) | Virus yield/cell (GC) |
| --- | --- | --- | --- | --- | --- |
| AAV1 | eGFP | 4.87E+12 | 100 | 4.87E+11 | 1.35E+04 |
| AAV2 | eGFP | 3.06E+11 | 150 | 4.59E+10 | 1.53E+03 |
| AAV5 | eGFP | 9.65E+12 | 150 | 1.45E+12 | 4.03E+04 |
| AAV6 | eGFP | 8.38E+11 | 200 | 1.68E+11 | 4.67E+03 |
| AAV8 | eGFP | 6.18E+13 | 100 | 6.18E+12 | 1.72E+05 |
| AAV9 | eGFP | 1.18E+13 | 250 | 2.95E+12 | 8.19E+04 |

AAV-hSyn-eGFP was packaged with serotypes 1, 2, 5, 6, 8, and 9 using the two-step purification method in small scale. Viral titers were determined by Q-PCR as described in Materials and Methods.

To test the purity of samples, comparable genome copies of each sample were resolved via SDS-PAGE and stained for protein content with silver stain (Fig. 2a). Similar amounts of virus for AAV1, 5, 8, and 9 (3E+10 to 1E+11 GC), and 2E+9 and 5E+9 GC for AAV2 and 6 (due to low titer and limited volume that could be loaded onto a well) were loaded onto the gel in Fig. 2a. Equal volumes of AAV preparations (3ul of each preparation) were loaded onto the gel in Fig. 2b. Two samples of AAV-hSyn-eGFP were purchased from commercial sources (Addgene.org and UPenn Vector Core) and resolved side-by-side on silver-stained gels as positive controls. The same AAV transfer plasmids for AAV2 and AAV9 were purchased from Addgene and UPenn Vector Cores and used for packaging AAVs using the two-step purification method. Silver-staining assay is typically used to detect the major AAV capsid proteins VP1, VP2, and VP3; and to compare the ratios of AAV capsid proteins to impurities. The amount of VP1, VP2, and VP3 is also indicative of the amount of AAV particles present in the preparation (Fig. 2b). The low genome copies of two-step purification for AAVs with serotypes 2 and 6 are evident on the gel in Fig 2b. However, AAV serotypes 1, 5, 8, and 9 had robust VP1, VP2, VP3 presentations. All two-step preparations had comparable purities to purchased commercial AAV preparations (Fig. 2a).

**Fig. 2.**
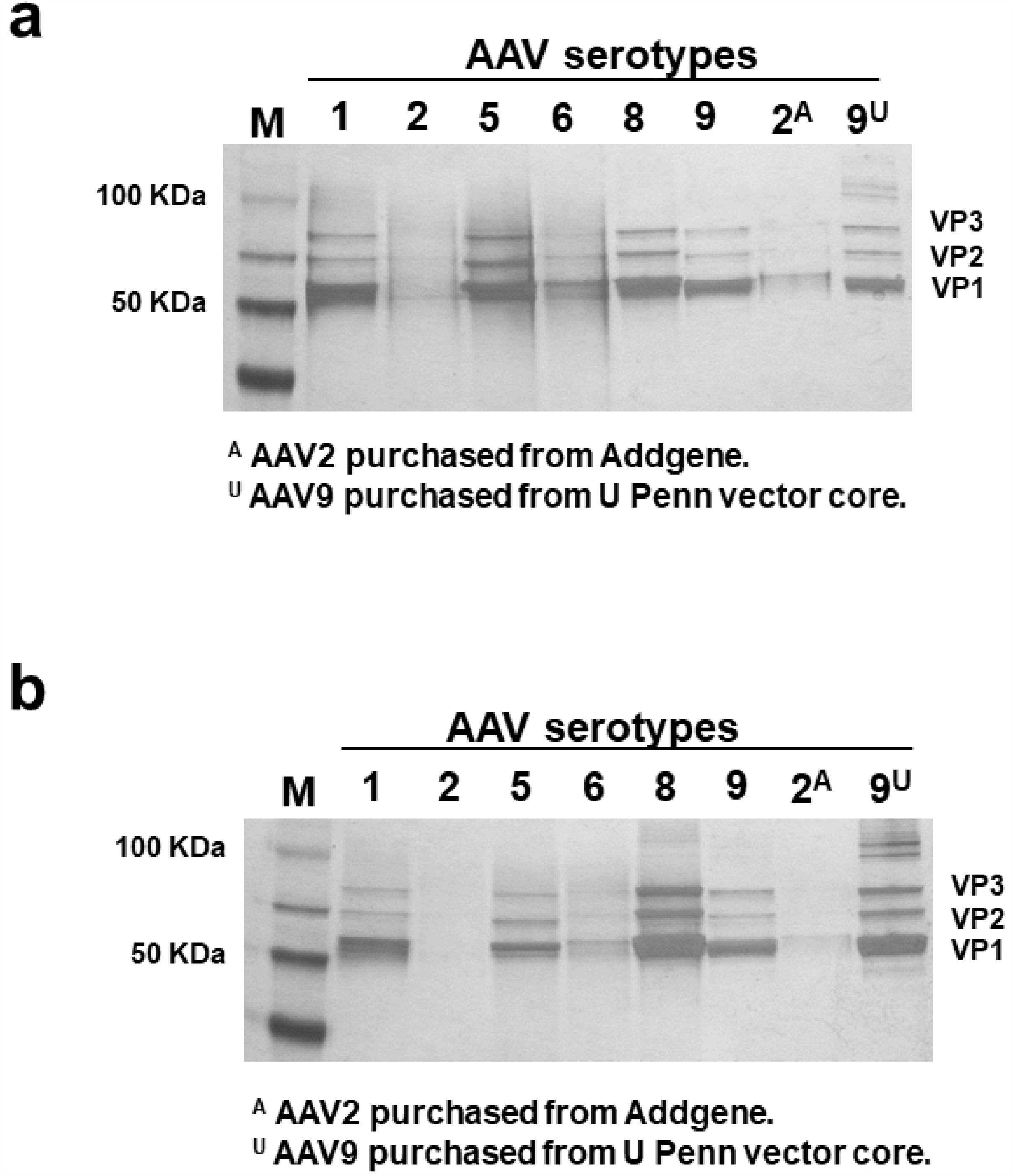
Comparison between purity and yield of AAV serotypes.

AAV-hSyn-eGFP was packaged with serotypes 1, 2, 5, 6, 8, and 9 using the two-step purification method. a. Similar amounts of virus for AAV1, 5, 8, 9, using the two-step purification and AAV2 and 9 purchased from Addgene and UPenn Vector Core (3E+10 to 1E+11 GC) were loaded onto the gel. Due to low titer and limited volume that could be loaded onto a well, only 2E+9 and 5E+9 GC of AAV2 and 6 purified with the two-step purification were resolved. Three major protein bands VP1, VP2, and VP3 are the most abundant proteins resolved by silver-staining. b. Equal volumes of AAV preparations (3ul) were loaded onto the gel and silver-stained. Two samples of AAV-hSyn-eGFP were purchased from commercial sources (Addgene.org and UPenn Vector Core) and resolved side-by-side on silver-stained gels as positive controls. Same AAV transfer plasmids for AAV2 and AAV9 were purchased from Addgene and UPenn Vector Cores and used for packaging AAVs in the two-step purification method.

Transmission electron microscopy (TEM) was used to visualize the structural features of the AAV particles and sample impurities. AAV9 sample prepared by the two-step purification method and its commercially purchased equivalent AAV9 from the UPenn Core were imaged using TEM (Fig. 3). As shown in the negative stain images, both samples have similarly stained particles and impurities. The purchased AAV9 from UPenn contained more particles and higher titer.

**Fig. 3.**
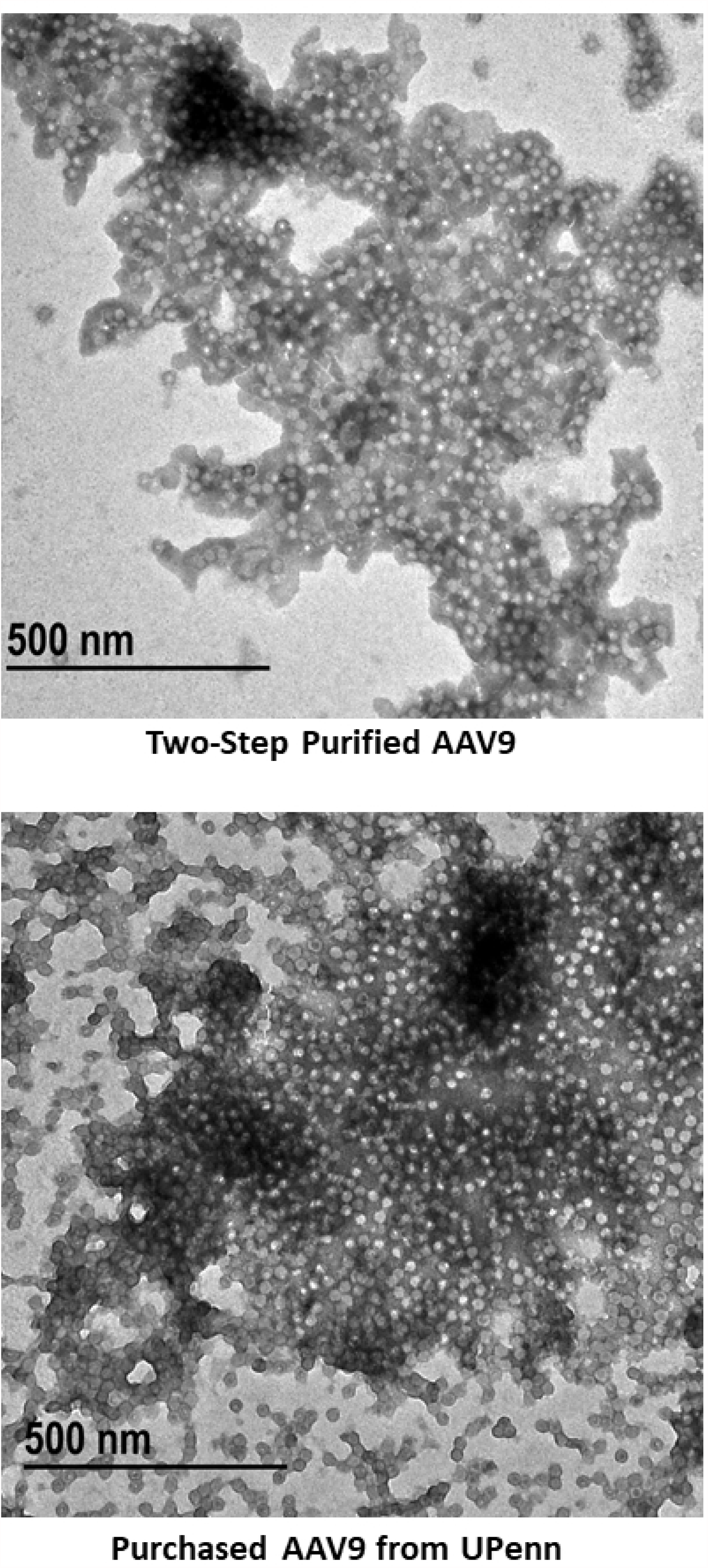
Transmission Electron Microscopy (TEM) images of AAV samples.

Equal volumes (5 μL) of AAV9 sample preparation by the two-step purification method and the AAV9 samples purchased from the UPenn Core were imaged by TEM as described in Materials and Methods. Negative stained images were obtained at 68000x.

### Ex-vivo AAV transduction

In addition to determining the genome copies for each AAV preparation and resolving the samples on silver-stained gels, we tested the transduction efficiency of each sample in organotypic mouse brain slice cultures. Human synapsin promoter is a robust promoter in central nervous system neurons and provides an effective way of comparing GFP expression in AAV transduced living tissue. Many small-scale AAV preparations are used for gene delivery to animal tissue in research laboratories [25,26].

Equal amounts of viral preparations were added dropwise on top of cultured brain tissues and monitored for expression of GFP by confocal microscopy. AAV expression was detected one week after infection. AAV preparations for serotypes 1, 5, 6, 8, and 9 transduced mouse neurons effectively (Fig. 4). Despite low titer, the AAV sample with serotype 6 transduced neurons as effectively as other serotypes. AAV with serotype 2, had the lowest yield and expressed the lowest levels of GFP. Positive control samples of AAV-hSyn-eGFP with serotypes 2 and 9 from commercial sources (Addgene.org and UPenn Vector Core, respectively) had comparable robust expressions similar to the two-step purified AAV preparations. The differences in GFP expression, and hence transduction, could also be due to tropism of different serotypes.

**Fig. 4.**
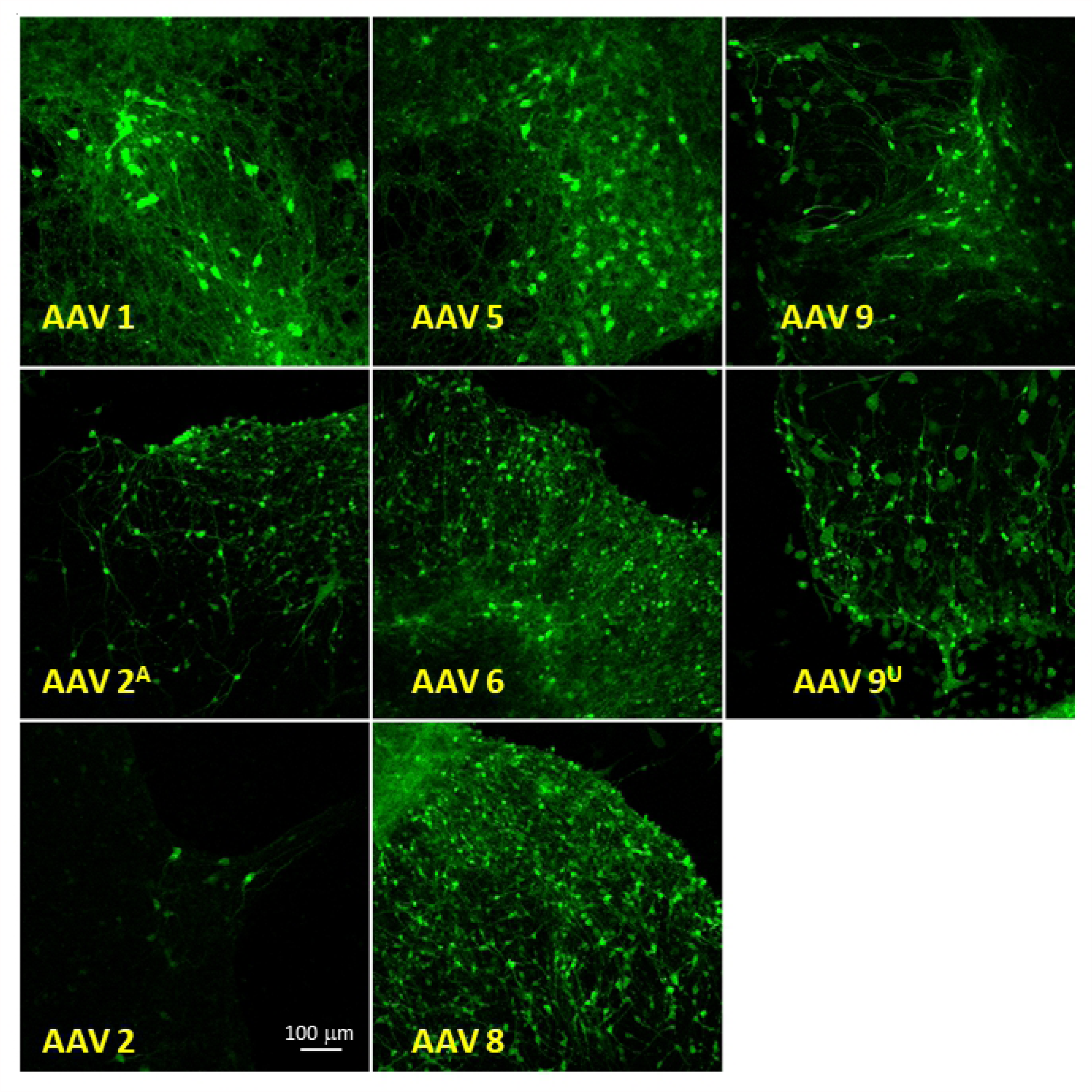
AAV transduction of cultured mouse brain slices.

Organotypic mouse brain slice cultures were prepared and transduced with various AAV serotypes. Confocal images depict GFP expression in cultured mouse neurons after two weeks. Despite low yield, AAV6 transduced cells as efficiently as higher yield AAV preparations. Commercially purchased samples were used as positive controls.

All two-step purified AAV samples, except for serotype 2, demonstrate effective transduction efficiencies with no observed toxicity. GFP expression in neurons was consistently monitored for 4 weeks with no observed changes in expression.

## Discussion

Rapid expansion of the AAV toolbox has availed a wide range of vectors with designer capsids, promoters, and/or selectable markers to researchers [8,25,27]. AAVs are preferred vectors for many *in vivo* gene transfer applications to non-dividing cells. In order to facilitate gene delivery for basic and clinical research, manufacturing practices for AAV production have been optimized to provide research and clinical grade preparations [10]. A myriad of AAV samples are commercially packaged and available for purchase. However, preparation of AAVs from customized vectors carrying genes with single nucleotide polymorphisms, dominant negative mutations, chimeric proteins, etc. are often costly and time consuming to prepare. Here, we describe a method to purify several AAV serotypes rapidly, economically, and in small-scale.

In this method, cell transfection and recovery of AAV particles from cell media is performed according to standard protocols. The virus is collected from the culture medium that has been incubated for five days as compared to traditional method of collecting AAV particles from freeze-thawed cell lysate in three days. During the five-day incubation following transfection, AAV particles are released and accumulated in media devoid of serum. Therefore, less impurities are present in the starting material compared to freeze-thawed cell lysates. The purification steps are simplified and shortened without jeopardizing quality. In this method, since AAVs are not purified through multiple steps, less sample is lost during purification steps, and less starting material is required. This protocol allows research laboratories to rapidly produce multiple AAV preparations from small amounts of starting material.

The purified virus stocks using the two-step method may also contain empty AAV viral capsids. The effect of empty capsids on transduction outcome is still unclear. The presence of empty capsids may increase the unwanted immune response; yet, some studies report that the existence of empty capsids is beneficial and increases transduction [28,29].

The standard purification protocols also rely on collecting AAV fractions from gradients and examination of each individual fraction for quality and quantity. Samples purified using cesium chloride also require an additional dialysis step to remove toxic cesium chloride. In this protocol, no toxic material is introduced into the collected samples and a protein concentrator device is used as a filtration device to remove impurities. The AAV samples prepared by the two-step purification method were resolved by SDS-PAGE, silver-stained, and imaged by TEM to demonstrate the purity of samples as compared to similar commercially obtained AAV preparations. The simplification of the purification protocol saves resources and time. The materials used in the two-step purification protocol are inexpensive, and the equipment used, such as centrifuges, are readily available in most research laboratories. This method also reduces the amount of generated hazardous waste.

Using this two-step purification protocol, we have prepared and tested AAVs with serotypes 1, 2, 5, 6, 8, and 9. Low recovery yields were observed for serotypes 2 and 6. Furthermore, serotype 2 preparation showed low levels of neuronal transduction in organotypic mouse brain slice cultures. Low yield may be due to binding of surface proteins of AAV serotype 2 to cell surface heparan sulfate proteoglycan (HSPG) that has been shown to reduce the number of AAV2 particles released into the culture media during virus production [30]. Each chimeric/designer AAV serotype should be assessed and validated individually for yield and transduction efficiency after purification with the described two-step purification method.

The two-step AAV purification method is an effective and easy-to-use method for research laboratories to produce small-scale, yet high-quality, AAV preparations. This technique enables researchers to rapidly create and test new AAV constructs in animal models and can be scaled up for bulk preparations.

## Acknowledgements

This research was supported by the Intramural Research Program of the National Institute of Health (NIH), National Institute of Environmental Health and Sciences (NIEHS). We are grateful to Dr. Carl Bortner and Dr. Jesse Cushman for critical reading of the manuscript and helpful advice. We would also like to acknowledge and thank Dr. Jerrel Yakel, Dr. Zhenglin Gu, Ms. Pattie Lamb, Ms. Deloris Sutton, and Dr. Bernd Gloss for their intellectual and technical contributions.

## Author Contribution

NPM, AP, SHC, and MW developed the concept of the protocol and performed experiments. ES provided the confocal imaging data. RDK provided the TEM images. NPM wrote the manuscript. All authors contributed to editing of the paper.

## Compliance with Ethical Standards

### Funding

This research was supported by the Intramural Research Program of the National Institute of Health (NIH), National Institute of Environmental Health and Sciences (NIEHS).

### Ethical Approval

Animals used in this study were ordered from Charles River and Jackson Laboratories, USA. All animal procedures complied with the institutional guidelines, NIH/NIEHS, animal care guidelines and were approved by the Animal Care and Use Committee (ACUC) at the NIH/NIEHS, animal Protocol # 2012-0004.

